# Genetic and epigenetic convergence at *MOBP* links ALS and PSP

**DOI:** 10.64898/2026.03.25.714147

**Authors:** Katherine Fodder, Megha Murthy, Joshua Harvey, Ehsan Pishva, Katie Lunnon, Towfique Raj, Kurt W Farrell, Rohan de Silva, Jack Humphrey, Conceição Bettencourt

**Author notes:** Corresponding Authors: Jack Humphrey, PhD, Icahn School of Medicine at Mount Sinai 1 Gustave Levy Plaza New York 10029, Conceição Bettencourt, PhD, Department of Neurodegenerative Disease UCL Queen Square Institute of Neurology 1 Wakefield Street London WC1N 1PJ United Kingdom.

## Abstract

**Background:** Myelin associated oligodendrocyte basic protein (*MOBP*) is an abundantly expressed oligodendrocyte gene. Genetic variation at the *MOBP* locus has been associated with risk for multiple neurodegenerative disorders, including progressive supranuclear palsy (PSP), amyotrophic lateral sclerosis (ALS), and frontotemporal dementia (FTD). Epigenetically, *MOBP* promoter hypermethylation and reduced expression have been reported in multiple system atrophy (MSA). Although *MOBP* is thought to play a role in oligodendrocyte morphology and myelin structure, how genetic and epigenetic variation at this locus influences gene regulation and contributes to disease risk remains poorly understood across neurodegenerative disorders.

**Methods:** We assessed genetic overlap and colocalisation at the *MOBP* locus across multiple neurodegenerative diseases. Disease-associated signals were fine-mapped to prioritise putative causal variants, and the strongest signals were subsequently investigated using molecular quantitative trait loci and post-mortem brain tissue datasets.

**Results:** ALS and PSP showed strong genetic association at *MOBP*, with highly overlapping associated variants and strong evidence that both diseases share the same causal signal. Overlap with frontotemporal dementia was weaker. Methylation quantitative trait locus colocalisation identified cg15069948, located near an exon junction within *MOBP*, as strongly colocalising with the ALS and PSP genetic risk signals. In PSP brain tissue, carriers of the rs1768208-T risk allele showed lower methylation at cg15069948. We additionally identified evidence for an interaction between rs1768208 genotype and cg15069948 methylation associated with altered *MOBP* 3′ untranslated region architecture.

**Conclusions:** These findings identify a shared ALS/PSP genetic–epigenetic mechanism at *MOBP* that is distinct from promoter-associated dysregulation in MSA. Together, they position *MOBP* as a key oligodendrocyte regulatory locus with both shared and disea e-specific roles in neurodegeneration.

## Background

Although neurodegenerative diseases are traditionally defined as distinct entities, considerable overlap in clinical features and neuropathology is commonly observed (1). Consistent with this, genetic, transcriptomic, and epigenetic studies increasingly implicate shared molecular pathways across subsets of these disorders. Genome-wide association studies (GWAS) have revealed both shared and disease-specific risk loci. Defining where their genetic architectures converge or diverge is crucial for uncovering common pathways of neurodegeneration and identifying therapeutic targets with potential cross-disease relevance.

Myelin associated oligodendrocyte basic protein (MOBP), which is almost exclusively expressed in oligodendrocytes (2) is linked to multiple neurodegenerative diseases through distinct mechanisms. Although its exact function remains unclear, MOBP is the third most abundant CNS myelin protein and is thought to stabilise myelin and support oligodendrocyte morphological differentiation (3,4). Functional studies by Schäfer et al. demonstrated that *MOBP* knockdown reduces surface area of MBP-positive oligodendrocytes and impairs myelin-like membrane formation (5).

Genetic variation in *MOBP* is associated with the risk of multiple neurodegenerative diseases, underscoring its biological relevance. Risk-associated SNPs have been identified in progressive supranuclear palsy (PSP)(6–10), corticobasal degeneration (CBD)(10), frontotemporal dementia (FTD)(11), amyotrophic lateral sclerosis (ALS)(12), and in APOE-ε4–positive Alzheimer’s disease (AD) (13). One key variant, rs1768208-T, an intronic SNP in *MOBP*, is robustly associated with PSP and CBD risk and with white-matter degeneration and faster executive decline in bvFTD (14). Although its functional impact is not fully understood, the risk allele has been associated with increased *MOBP* expression in PSP brain tissue (15,16).

DNA methylation studies in multiple system atrophy (MSA) have identified *MOBP* as a key dysregulated locus. The *MOBP* promoter has been reported as the most differentially methylated region in MSA cerebellar white matter, with similar changes also observed in frontal and occipital white matter (17). The presence of these changes in mildly affected regions suggests they occur early in disease and may contribute to pathogenesis. *MOBP* expression was reduced in cerebellar white matter and in oligodendrocytes of MSA compared to controls (18). Follow-up analyses showed that *MOBP* promoter hypermethylation is inversely correlated with *MOBP* expression (2), consistent with DNA methylation driving *MOBP* downregulation. At the protein level, MOBP isoform abundance and ratios were unchanged in the white matter of MSA, whereas PSP showed increased expression of all three isoforms and a shift toward larger isoforms (2). MOBP shows disease-specific localisation in cerebellar white matter, restricted to myelin in controls but enriched in glial cytoplasmic inclusions in MSA, where it interacts with α-synuclein (2). Sequestration into Lewy bodies has also been reported in PD and LBD (19).

Given our previous work and the involvement of *MOBP* in common and rare neurodegenerative diseases, understanding shared mechanisms involving this gene in disease processes could give crucial insight into common pathways that lead to neurodegeneration and provide targets for therapeutic intervention in a broader range of diseases. Here, we investigate how *MOBP* is dysregulated across several neurodegenerative diseases through investigations of genetic, epigenetic and transcriptomic data. We identify a strong signal of shared genetic association between PSP and ALS prioritising the same causal SNPs, which is not observed in the other neurodegenerative diseases. DNA methylation analysis identified a CpG site (cg15069948) at the *MOBP* locus that colocalises with the PSP and ALS genetic signal, providing evidence for a shared regulatory mechanism linking genetic risk to epigenetic variation at this locus. Additionally, our work suggests that an interaction between DNA methylation levels and disease risk allele at this locus may alter splicing and the architecture of the 3’ untranslated region (UTR) at *MOBP*.

## Methods

### Datasets (GWAS, mQTL, and eQTL)

To investigate genetic architecture at the *MOBP* locus across neurodegenerative diseases, we performed colocalisation analyses using publicly available GWAS summary statistics. For each disease, we obtained full GWAS summary statistics and accompanying lists of genome-wide significant loci from the respective studies (Table 1). When required, missing fields were harmonised as follows: standard errors were derived from reported effect sizes and P-values; minor allele frequencies were supplemented using European reference data from the 1000 Genomes Project; and SNP coordinates or RS IDs were matched using Ensembl GRCh37/hg19 (20).

**Table 1.**
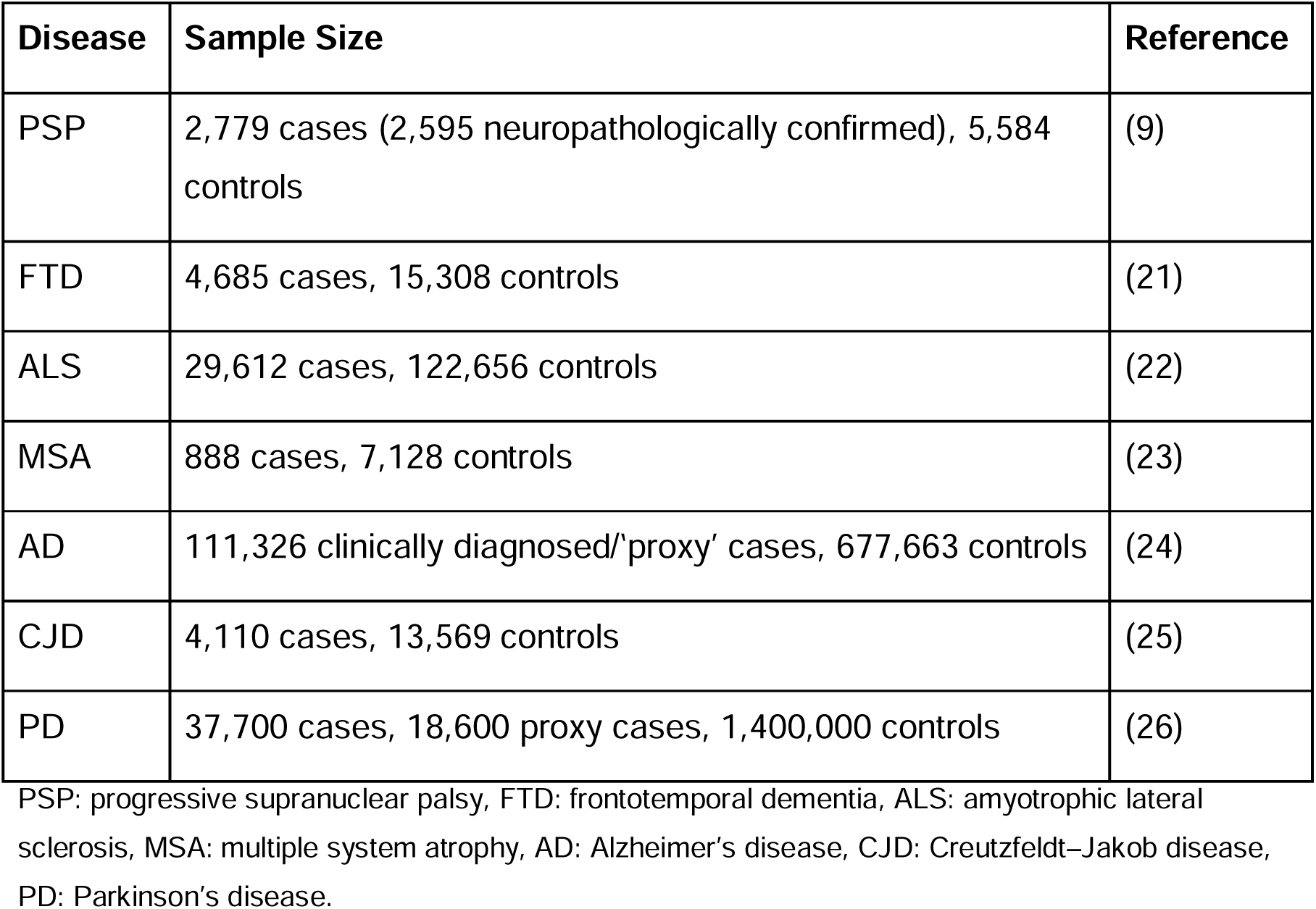
Description of the disease GWAS used to investigate genetic associations at *MOBP* across neurodegenerative diseases.

Summary statistics from human bulk brain methylation quantitative trait loci (mQTL), oligodendrocyte specific single-nucleotide expression quantitative trait loci (eQTL) and protein quantitative trait loci (pQTL) were made available by Cifello et al. (27) and accessed through the NIAGADS database (https://dss.niagads.org/datasets/ng00184/).

For all the analyses, we used a region comprised between 1 Mb upstream and downstream of the coordinates *of MOBP* (GRCh37 Chromosome 3: 39,508,689-39,570,970) to investigate association between disease SNPs, mQTLs, eQTLs and pQTLs.

### Colocalisation Analysis

As an initial exploratory step, we assessed the similarity of association patterns across neurodegenerative diseases and molecular traits at the *MOBP* locus using correlation-based approaches. First, we calculated pairwise correlations between SNP-level GWAS *p*-values across diseases within the *MOBP* region (±1 Mb around the gene) to evaluate the extent of shared genetic signal.

Next, using bulk brain mQTL summary statistics, we extracted data for all CpG sites within ±1 Mb of *MOBP* and, for each CpG, computed SNP-wise correlations between *p-values* for mQTL disease-associated GWAS SNPs across the locus. This analysis was used to identify CpGs whose genetic regulation showed concordant patterns with disease association.

We next performed formal colocalisation analysis using the COLOC R package (version 3.2-1) (28). COLOC evaluates whether two association signals, such as disease GWAS signals and mQTL or eQTL effects, are consistent with a shared causal variant within a defined genomic region. For each disease GWAS dataset, we extracted all SNPs within the *MOBP* locus (±1 Mb around the gene). Prior probabilities in COLOC reflect assumptions about the likelihood that a SNP is associated with trait 1 (p1), trait 2 (p2), or both traits (p12). As only a small proportion of SNPs are expected to be causal, default priors are typically low (p1 = p2 = 1×10^−^□; p12 = 1×10^−^□). However, given prior evidence supporting association at the *MOBP* locus, we increased these values (p1 = p2 = 1×10^−^³; p12 = 1×10^−^□) to reflect a higher prior probability of association and colocalisation in this targeted analysis.

Approximate Bayes Factors (ABFs) were computed internally by COLOC to estimate posterior support for each hypothesis. To complement the COLOC ABF framework and allow for multiple causal variants within a region, we applied the SuSiE (29) fine-mapping model to both the GWAS and QTL datasets. SuSiE identifies credible sets of putative causal SNPs by modelling associations as the sum of multiple independent effects, returning a posterior inclusion probability (PIP) for each SNP and one or more credible sets.

### Genotyping of *MOBP* rs1768208

Several SNPs within *MOBP* have been associated with neurodegenerative diseases, with rs1768208 repeatedly reported as a lead variant (6,7). We therefore selected rs1768208 (Chr3:39,508,968, GRCh37) for targeted genotyping in a subset of samples with available DNA methylation data.

A 381 bp PCR fragment encompassing the variant was amplified using flanking primers (forward: 5′-TCC TCT CAA GCC TCA AAC TCT C-3′; reverse: 5′-GGC AAC TCA GCC CAG AAA TTT G-3′). Each 15 μl PCR reaction contained 7.5 μl Promega GoTaq® Green Master Mix, 2.25 μl of each primer (3 μM), 2 μl genomic DNA (∼100 ng), and 2 μl dH₂O, and was amplified using a standard touchdown protocol. PCR products were verified by agarose gel electrophoresis and purified with an ExoSAP mix (exonuclease I and FastAP). Sanger sequencing was performed by Source Bioscience using the forward primer, and chromatograms were visualised in Benchling (https://benchling.com/) to determine rs1768208 genotypes.

To assess the effect of rs1768208 on DNA methylation, we analysed frontal lobe white-matter methylation data generated previously from control, MSA, PD, and PSP brain tissue (31) (**Supplementary Table S1**). Following standard pre-processing, quality control, and normalisation carried out by Murthy et al. (30), adjusted M-values, i.e. logit transformation of beta-values, were obtained for samples overlapping those successfully genotyped at rs1768208. These values were used to test for DNA methylation differences in relation to rs1768208 genotype (presence/absence of the risk allele (T)).

### Matched DNA methylation, gene expression and splice-junction analysis at *MOBP*

To investigate whether genetic and epigenetic variation at the *MOBP* locus was associated with *MOBP* expression changes and/or transcript processing, we analysed matched genotype, DNA methylation, short read RNA-sequencing (RNA-seq) gene expression and splice-junction data from post-mortem brain samples (31,32). A total of 88 samples derived from human prefrontal cortex post-mortem tissue were included where rs1768208 genotype, cg15069948 methylation, relevant phenotypic covariates and RNA-seq-derived expression or splice-junction measurements were available.

rs1768208 genotype was modelled as T-carrier status, comparing individuals carrying at least one T allele (TT or TC) with non-carriers (CC). DNA methylation at cg15069948 was adjusted prior to downstream modelling using a linear model adjusting for age, sex, methylation array plate and estimated oligodendrocyte proportion (NeuN-negative/SOX10-positive fraction)(33). Adjusted methylation values were calculated as model residuals plus the intercept and used in subsequent association analyses.

To assess whether genetic and epigenetic variation influenced total *MOBP* expression, linear models were used to test the association of log_2_-transformed *MOBP* expression with rs1768208 genotype, adjusted cg15069948 methylation and their interaction, adjusting for oligodendrocyte proportion, age, RNA integrity number (RIN) and sex.

*MOBP* splice junction usage was quantified using the Leafcutter (34) workflow and software. Raw RNA sequencing reads (Illumina NovaSeq 6000) underwent initial quality control profiling using FastQC, with automated trimming applied to strip sequencing adapters and eliminate low-quality terminal bases. The remaining high-quality reads were aligned to a prebuilt human reference genome index (GRCh38 assembly, generated with STAR function genomeGenerate) using the splice-aware STAR aligner (Version 2.7.11, twopassMode = Basic, outSAMstrandField = intronMotif). Resulting bam files were converted to junction file format and reads aligned to chromosome three were extracted using samtools and regtools. Intron clustering was conducted using the leafcutter_cluster_regtools python script with settings of 50 split reads supporting each cluster and allowing introns of up to 500kb. For each LeafCutter cluster, raw intron counts were converted to proportional junction usage by dividing each junction count by the total count for all junctions within the same cluster in each sample. Junctions were filtered to retain those with sufficient read support, defined as mean cluster read depth ≥20 and total junction read count ≥20 across samples. For each retained junction, raw proportional usage was tested for association with rs1768208 genotype, cg15069948 methylation and their interaction using linear regression. Models included oligodendrocyte proportion, age, RNA integrity number and sex as covariates. Separate models tested genotype-only effects, methylation-only effects and genotype-by-methylation interactions. For consistency with GWAS summary statistics, coordinates were converted to GRCh37 for visualisation.

A schematic overview of the methodological framework is shown in Figure 1.

**Fig. 1.**
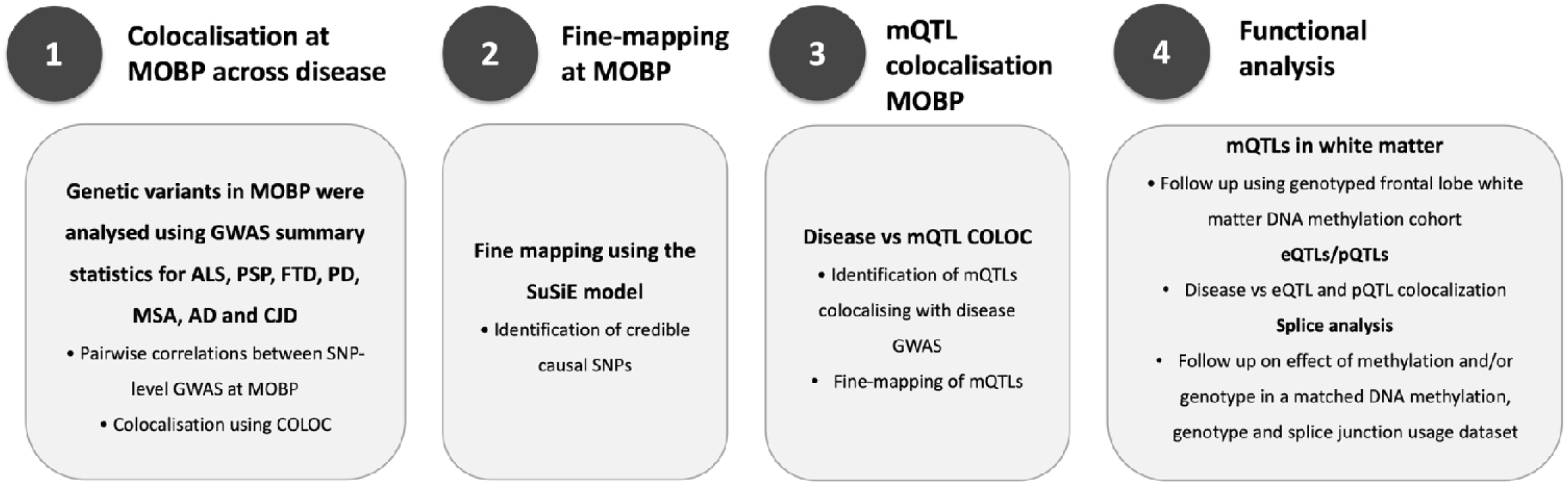
Schematic overview of the analytical workflow used to investigate the *MOBP* locus across neurodegenerative diseases. 1) Cross-disease colocalisation analysis at the *MOBP* locus using GWAS summary statistics from ALS, PSP, FTD, PD, MSA, AD and CJD, including pairwise SNP-level comparisons and Bayesian colocalisation (COLOC). 2) Fine-mapping of the *MOBP* locus using the SuSiE model to identify credible sets of putative causal variants. 3) Colocalisation of disease-associated variants with brain DNA methylation quantitative trait loci (mQTLs), including fine-mapping of disease-colocalising mQTL signals. 4) Functional follow-up analyses of prioritised variants, including validation of mQTLs in a genotyped frontal white matter DNA methylation cohort, disease versus eQTL/pQTL colocalisation analyses, and investigation of the effects of genotype and DNA methylation on *MOBP* splice junction usage in matched datasets derived from post-mortem brain tissue. MOBP, myelin-associated oligodendrocyte basic protein; ALS, amyotrophic lateral sclerosis; PSP, progressive supranuclear palsy; FTD, frontotemporal dementia; AD, Alzheimer’s disease; MSA, multiple system atrophy; PD, Parkinson’s disease; CJD, Creutzfeldt-Jakob disease; GWAS, genome-wide association study; SNP, single-nucleotide polymorphism; COLOC, Bayesian colocalisation analysis; SuSiE, Sum of Single Effects; mQTL, methylation quantitative trait loci; eQTL, expression quantitative trait loci; pQTL, protein quantitative trait loci.

## Results

### Colocalisation analysis reveals shared *MOBP* genetic risk variants between ALS and PSP

To explore the genetic landscape surrounding *MOBP* across multiple neurodegenerative diseases, we examined a ±1 Mb window around the locus using summary statistics from seven GWAS (**Table 1**). Only ALS and PSP exhibited a clear, highly significant association peak at *MOBP* (**Figure 2A, Supplementary Figure 1A**), with several SNPs reaching genome-wide significance (P < 5 × 10⁻□). FTD showed several variants reaching suggestive significance (P < 1 × 10⁻□) but none passing the genome-wide significance threshold, possibly reflecting a smaller sample size and FTD clinicopathological heterogeneity. None of the other diseases displayed evidence of genetic association at this locus.

**Fig. 2.**
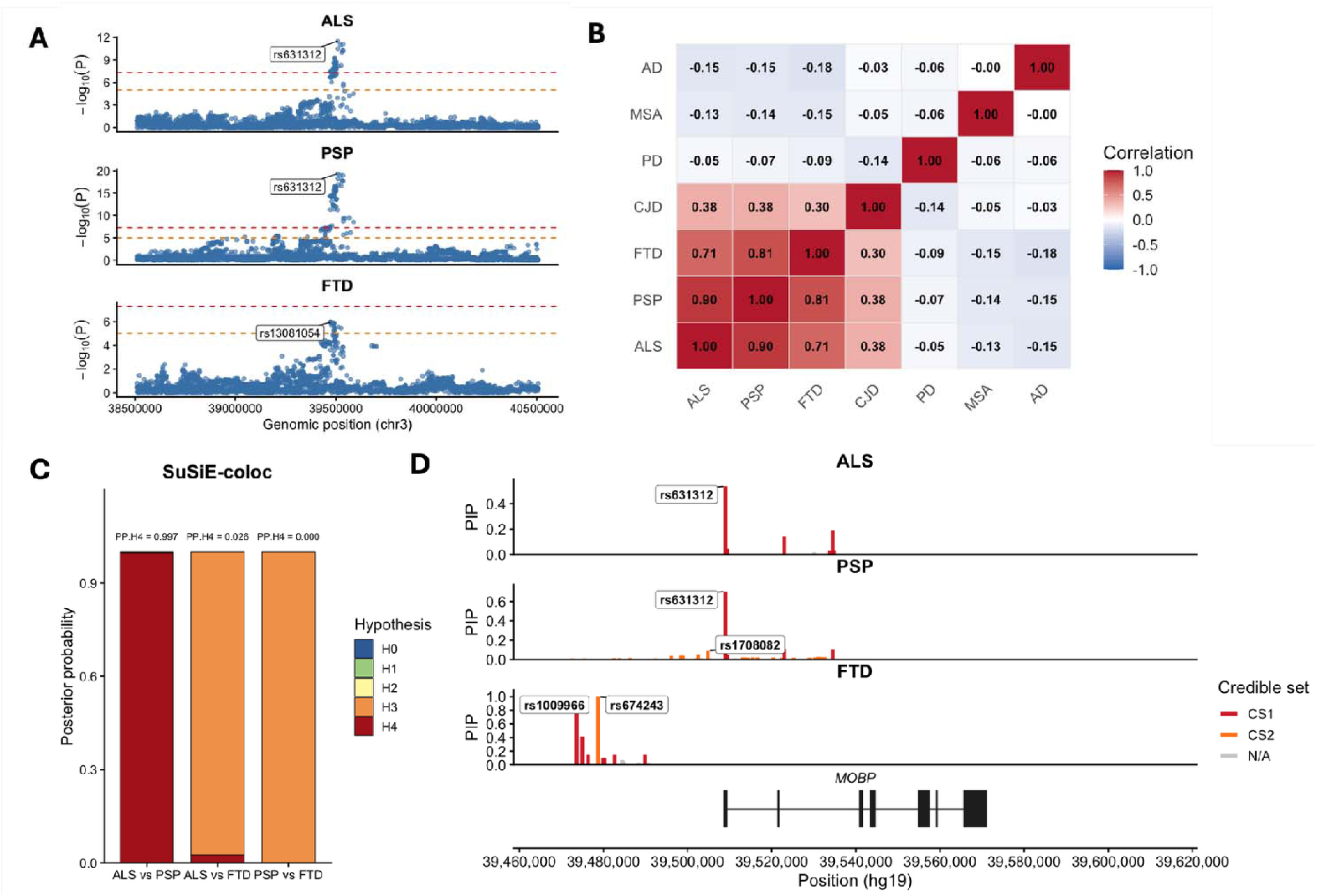
Genetic architecture and cross-disease colocalisation of the *MOBP* locus across neurodegenerative diseases. A) Regional Manhattan plots for ALS, PSP and FTD across a ±1 Mb window around the *MOBP* locus. Points represent single SNP associations, plotted as –log_10_(P). Red and orange dashed lines indicate genome-wide significance (P = 5×10⁻□) and suggestive significance (P = 1×10⁻□), respectively. B) Pairwise correlation heatmap of GWAS –log_10_(P) values across the *MOBP* region in a range of diseases. C) SuSiE-coloc posterior probabilities for disease-pair comparisons at *MOBP.* D) Regional fine-mapping plots showing SuSiE-derived credible sets for ALS, PSP, and FTD across the canonical *MOBP* locus. SNPs are coloured by credible set membership: CS1 (primary credible set), CS2 (secondary credible set), or non-credible set (N/A). ALS, amyotrophic lateral sclerosis; PSP, progressive supranuclear palsy; FTD, frontotemporal dementia; AD, Alzheimer’s disease; MSA, multiple system atrophy; PD, Parkinson’s disease; CJD, Creutzfeldt-Jakob disease; SNP, single-nucleotide polymorphism; PP, posterior probability.

To determine whether different diseases share the same underlying *MOBP* risk-associated variant(s), we next quantified pairwise similarity across all SNPs in the region. ALS, PSP, and FTD showed strong positive pairwise correlations (Pearson r = 0.70–0.90, p < 1 ×10^−200^), suggesting substantial overlap in disease-associated genetic architecture at the *MOBP* locus (**Figure 2B**). CJD displayed moderate correlation with these three diseases (r = 0.30–0.38, p < 0.001), indicating a weaker but detectable shared component. All other disease pairs showed weak or no correlation, consistent with the absence of significant signal at this locus. Together, these results demonstrate that ALS and PSP share the most pronounced and consistent genetic signal at *MOBP*, with FTD potentially showing a weaker overlap.

Given that ALS, PSP and FTD were the only diseases showing notable genetic associations at the *MOBP* locus, we prioritised detailed investigation of their relationships. We used the COLOC framework to quantify whether pairs of diseases shared the same causal genetic variant at *MOBP*. We observed very strong evidence of genetic colocalisation between ALS and PSP, with PP.H4 = 0.997 (PP.H4: both diseases are associated and share a single causal variant) (**Supplementary Figure 1B**), indicating that these diseases almost certainly share the same causal variant at *MOBP*. In contrast, ALS vs FTD and PSP vs FTD showed minimal support for colocalisation (PP.H4 = 0. 28 and 0.31, respectively) (**Supplementary Figure 1B**). For both latter comparisons, most of the posterior probability was assigned to PP.H3, suggesting that although FTD exhibits a signal in this region, its causal variant(s) are distinct from those driving ALS and PSP risk (PP.H3 = 0.69 and 0.68 for FTD vs ALS and PSP, respectively). SNP-level posterior probabilities identified a small number of variants with high PP.H4 that coincided with the association peak in both ALS and PSP, consistent with a shared signal at the *MOBP* locus (**Figure 2B**). In contrast, for FTD, SNPs with higher PP.H4 did not correspond to the top FTD-associated variants (**Supplementary Figure 1C**), supporting a distinct underlying signal. These results indicate that *MOBP* harbours a shared ALS–PSP genetic risk mechanism, whereas FTD shows a genetically separate architecture within the same locus.

As GWAS often identifies broad association peaks containing many SNPs in linkage disequilibrium (LD), fine-mapping methods are required to resolve these peaks and identify which specific variants are most likely to be causal. By applying SuSiE fine-mapping to ALS, PSP, and FTD summary statistics, we aimed to identify the likely causal SNP(s) driving association in each disease and determine whether ALS and PSP share the same causal SNP(s), as well as establishing whether the FTD signal is indeed independent. We applied COLOC.SuSiE to assess colocalisation using fine-mapped posterior probabilities rather than single-SNP signals (**Figure 2C**). The results mirrored and strengthened the COLOC.ABF findings, with high PP.H4 for ALS-PSP (0.99) and low PP.H4 for ALS-FTD and PSP-FTD (PP.H4 = 5.3 × 10^−^□ and 1.7 × 10^−^¹□, respectively). PSP resolved into two credible sets (**Figure 2D**), with a primary CS1 centred on rs631312 and a smaller secondary CS2. ALS showed a single credible set (CS1) comprising six SNPs, four of which overlapped with PSP CS1 (rs631312, rs1768208, rs616147, rs1768190), supporting a shared fine-mapped signal. The lead SNP in both diseases was rs631312 (chr3:39,508,968, GRCh37), which had the highest posterior inclusion probability (PSP PIP = 0.70; ALS PIP = 0.54). In contrast, FTD produced a single, non-overlapping credible set located upstream of the ALS/PSP signal (top SNP chr3:39,473,59, GRCh37), supporting an independent genetic association at the *MOBP* locus (**Figure 2D**).

### Genetic associations with *MOBP* methylation links ALS and PSP

Given our previous finding of aberrant DNA methylation at the *MOBP* locus in neurodegeneration (2,17), we next investigated whether genetic variants associated with ALS, PSP, or FTD colocalise with methylation quantitative trait loci (mQTLs) in human brain tissue (27) (N = 636, Dorsolateral prefrontal cortex). This allowed us to identify methylation sites (CpGs) within *MOBP* whose methylation levels are most strongly associated with the genetic risk variants. As with the GWAS–GWAS comparisons, we quantified the similarity of association signals across all SNPs in the region between GWAS and mQTL datasets. mQTLs for several CpGs showed strong positive P-value correlations with disease GWAS signals. The strongest correlations between ALS and PSP GWAS signals and brain mQTLs were observed at cg15069948 and cg07405330, both located within the *MOBP* gene body (cg15069948: Pearson r = 0.83 and 0.86, p < 1×10^−200^; cg07405330: Pearson r = 0.74 and 0.80, p < 1×10^−200^ for ALS and PSP, respectively) (**Supplementary Table S2**). At FTD, distinct CpGs showed the highest correlation with disease, with the top signal at cg06598794 (Pearson r = 0.82, p < 1×10^−200^), and highly correlated CpGs were predominantly located in the *MOBP* promoter region (**Supplementary Figure 2A**).

We then applied COLOC.ABF to test for colocalisation between mQTLs at all CpG sites and GWAS signals for each of the three diseases. This revealed a small number of CpGs showing high PP.H4 (shared causal variant between methylation changes and disease risk) (**Supplementary Table S3**), with cg15069948 emerging as the strongest candidate across ALS and PSP (PP.H4 = 0.982 for both comparisons, **Figure 3A**). This indicates a 98% probability that ALS and PSP share a single causal variant with the mQTL controlling methylation at cg15069948. We observed only modest colocalisation between this CpG and the FTD GWAS (PP.H4 = 0.43, **Figure 3A**). Although several CpGs showed strong pairwise correlation between FTD GWAS and mQTL signals, none demonstrated strong evidence of colocalisation (**Supplementary Table S3**). The risk allele (G) of the fine-mapped lead disease SNP for ALS and PSP (rs631312), was associated with decreased methylation at both top correlated CpG sites (cg15069948 and cg07405330, both in *MOBP* gene body) (**Figure 3B**). We further evaluated the rs631312-G mQTL effect for cg15069948 across two additional brain mQTL datasets from distinct brain regions, observing concordant negative effects; Parietal cortex (β = −0.0119, p = 7.18 × 10⁻□, N = 419), and Parahippocampal gyrus (β = −0.00647, p = 0.0102, N = 194)(27), supporting a reproducible association between the ALS/PSP risk allele and reduced *MOBP* gene-body methylation.

**Fig. 3.**
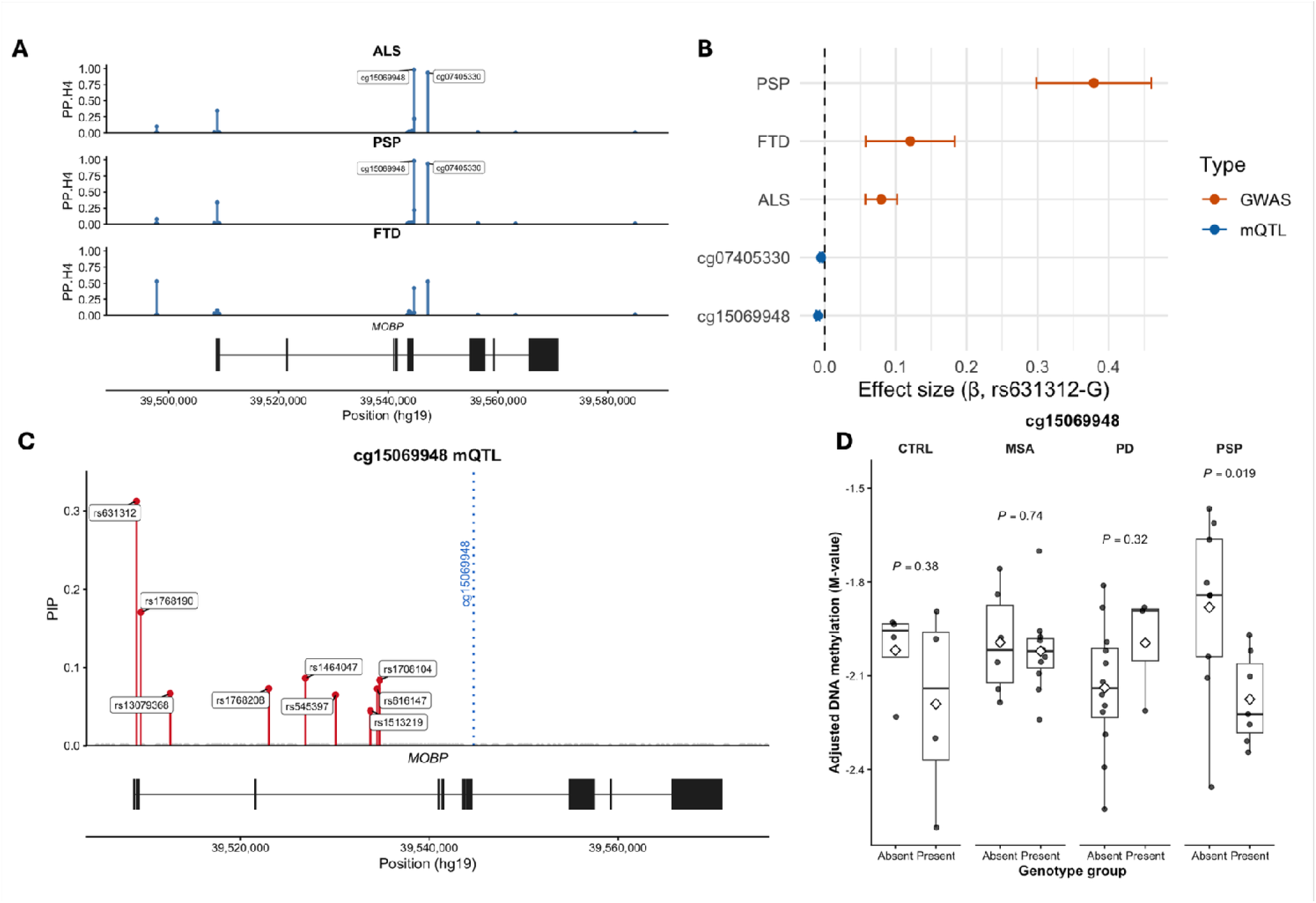
Genetic-epigenetic colocalisation and fine-mapping of mQTLs at the *MOBP* locus. A) DNA methylation probes showing the strongest evidence of colocalisation between *MOBP* mQTLs and disease GWAS signals across neurodegenerative disorders. Points/bars show posterior probability for the shared causal variant model B) Effect size estimates (β ± 95% CI) for rs631312 across disease GWAS and ROSMAP *MOBP* mQTL datasets, harmonised to the risk G allele. C) Fine-mapping of the *MOBP* mQTL signal, showing posterior inclusion probabilities for variants within the credible set. D) Frontal lobe white matter DNA methylation levels at cg15069948 across neurodegenerative disease groups, stratified by SNP risk allele presence. Differences in methylation between genotype groups were assessed using two-sample t-tests within each disease group. Boxplots show delta M-values for control, MSA, PD and PSP samples, with statistical comparisons (p-values) indicated. ALS, amyotrophic lateral sclerosis; PSP, progressive supranuclear palsy; FTD, frontotemporal dementia; MSA, multiple system atrophy; PD, Parkinson’s disease; SNP, single-nucleotide polymorphism; PP, posterior probability; mQTL, methylation quantitative trait loci; CS, credible set.

Fine-mapping of the cg15069948 mQTL revealed a single credible set (CS1) centred on the *MOBP* locus (**Figure 3C**). CS1 contained nine SNPs, with rs631312 showing the highest inclusion probability (PIP = 0.33) (**Figure 4C**). To assess whether the mQTL credible set overlaps biologically with disease-associated SNPs, we compared the SuSiE-derived mQTL CS1 with the fine-mapped credible sets from ALS, PSP, and FTD. All mQTL CS1 were present within either/both ALS and PSP CS1 sets. These shared variants include rs631312, rs1768208, rs1768190 and rs616147, the SNPs that also drive the strongest GWAS associations in both diseases.

**Fig. 4.**
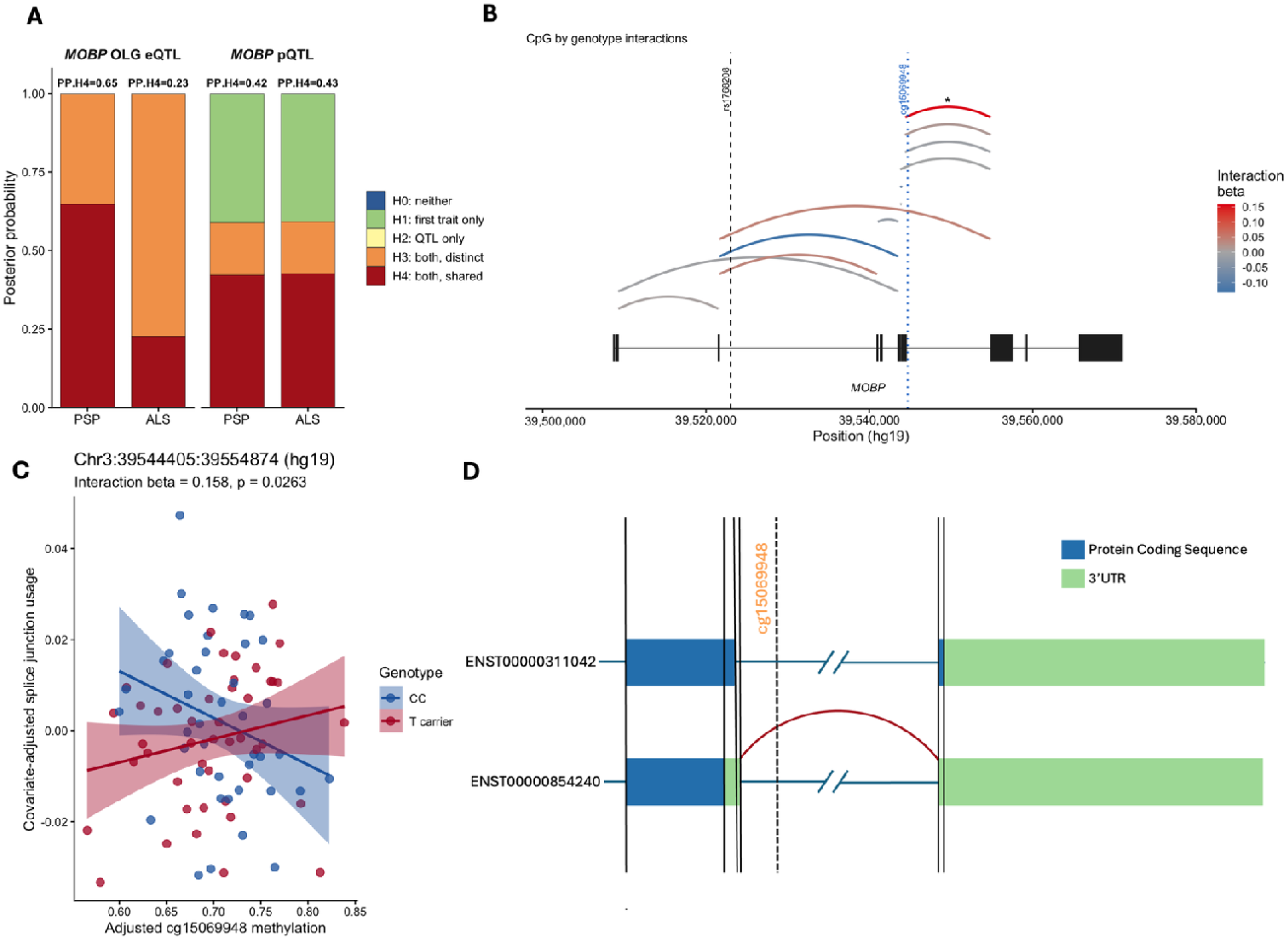
Genetic and epigenetic variation at the *MOBP* locus is associated with altered transcript processing near the 3′UTR transition. A) COLOC.ABF posterior probabilities for colocalisation between disease GWAS signals and *MOBP* oligodendrocyte-specific eQTLs and brain pQTLs. Stacked bars show posterior support for each colocalisation hypothesis, with PP.H4 values indicating evidence for a shared causal variant between disease risk and the molecular QTL signal. B) Schematic of *MOBP* cluster 890 splice junctions showing genotype-by-methylation interaction effects. Splice junctions are coloured according to the estimated interaction beta, with red indicating a positive interaction and blue indicating a negative interaction. Significance (p < 0.05) is indicated by an asterisk. The positions of rs1768208 and cg15069948 are shown relative to the *MOBP* gene structure. C) Interaction plot for the top splice junction, showing the relationship between adjusted cg15069948 DNA methylation and covariate-adjusted splice junction usage stratified by rs1768208 genotype group. Lines represent linear regression fits with 95% confidence intervals for CC individuals and T-allele carriers. The estimated interaction coefficient and corresponding P-value are shown. D) Schematic representation of selected *MOBP* transcript structures around the alternative 3′UTR transition. Protein-coding sequence and 3′UTR regions are shown separately, highlighting differences in transcript architecture near cg15069948 (Not to scale). *MOBP*, myelin-associated oligodendrocyte basic protein; ALS, amyotrophic lateral sclerosis; PSP, progressive supranuclear palsy; eQTL, expression quantitative trait locus; pQTL, protein quantitative trait locus; GWAS, genome-wide association study; SNP, single-nucleotide polymorphism; PP, posterior probability.

Whilst COLOC.ABF already indicated strong support for a shared causal variant (PP.H4 = 0.99 for both ALS–mQTL and PSP–mQTL comparisons), COLOC.SuSiE provided even stronger evidence by demonstrating that the same credible set accounts for both the mQTL signal and the ALS/PSP GWAS signals (**Supplementary Figure 2B**).

In contrast, FTD showed no credible-set overlap with the mQTL (PP.H4 = 0.16; zero shared SNPs), with posterior support of both traits being associated with different causal variants.

To complement the analysis of genetic variation at *MOBP* and any associated DNA methylation variation across neurodegenerative diseases, we undertook genotyping of the SNP rs1768208 (which is in high LD with the lead SNP rs631312, r^2^ = 0.9491 in European populations) to investigate any effects of genotype on DNA methylation in samples (N= 62) we had previously generated DNA methylation profiles of frontal lobe white matter across PSP, MSA and PD (30). The same CpG identified above as being an mQTL with strong association with PSP and ALS (cg15069948) showed nominally significant decrease in methylation within PSP samples with presence of the risk allele (P = 0.019) (**Figure 3D**), but not in the other neurodegenerative diseases. The direction of effect, a decrease in DNA methylation being associated with the presence of the risk allele (rs1768208-T), is in accordance with the direction of effect from the mQTL summary statistics (**Figure 3B**).

### Functional evaluation of genetic and epigenetic variation suggests a genotype by methylation interaction influencing *MOBP* alternative splicing

To further investigate the potential functional consequences of genetic variation at the *MOBP* locus, we tested whether disease-associated GWAS signals colocalised with oligodendrocyte-specific single-nucleus eQTLs (N = 736, Dorsolateral prefrontal cortex) or brain pQTLs (N = 416, Dorsolateral prefrontal cortex)(27) (**Figure 4A**). Colocalisation with the oligodendrocyte eQTL signal was inconsistent across diseases, with moderate support for a shared causal variant with PSP GWAS (PP.H4 = 0.65), but weaker support for ALS (PP.H4 = 0.23). In ALS, posterior probability was instead predominantly assigned to the H3 model, consistent with distinct disease and eQTL association signals at the locus. These findings suggest that altered total *MOBP* expression in oligodendrocytes is unlikely to fully account for the disease-associated signal. In contrast, *MOBP* pQTLs showed modest and comparable evidence of colocalisation with both PSP and ALS GWAS (PP.H4 = 0.42 and 0.43, respectively), but posterior support remained distributed across competing models. Therefore, while genetic variation at this locus may influence *MOBP* protein abundance, the pQTL results also do not provide strong evidence for a single shared causal variant with disease risk. Notably, these eQTL and pQTL colocalisation signals were substantially weaker than the mQTL colocalisation observed with PSP and ALS, indicating that the strongest molecular evidence at this locus was observed for DNA methylation rather than *MOBP* overall expression or protein abundance.

Given the strong evidence that disease-associated genetic variation at the *MOBP* locus colocalised with DNA methylation at cg15069948, we next utilised a bulk brain tissue dataset (N = 88, prefrontal cortex) with matched genotype, DNA methylation and RNA sequencing data to investigate whether changes in gene expression at *MOBP* could be due to interactions between DNA methylation and genotype. Consistent with the summary statistics eQTL analyses, we found no evidence that rs1768208 genotype influenced overall *MOBP* expression levels (beta = 0.01, p = 0.97) and saw no association between DNA methylation levels at cg15069948 and overall *MOBP* expression (beta = -1.14, p = 0.64) (**Supplementary Table S4**). We next tested for an interaction between genotype and methylation at cg15069948, and again, found no such effects on total *MOBP* expression (interaction beta = -0.63, p = 0.90) (**Supplementary Table S4**).

Given our previous observation suggesting that *MOBP* protein expression and isoform proportions are altered in PSP, with a shift towards higher molecular weight isoforms (2), we next investigated whether genetic and epigenetic variation at the *MOBP* locus might influence transcript processing rather than total *MOBP* expression. No individual splice junction showed a significant association with genotype or methylation alone (**Supplementary Table 5**). However, interaction analysis identified the strongest genotype-by-methylation interaction at a junction spanning the alternative 3′ region of *MOBP* (Chr3:39544405:39554874, GRCh37, β = 0.158, p = 0.0263; **Figure 4B,C, Supplementary Figure 3**). Notably, cg15069948 lies within the genomic interval spanned by this junction. The interaction-associated junction connects an upstream exon containing a short annotated 3′UTR segment to a downstream 3′UTR-containing exon, whereas the alternative transcript structure enters the 3′UTR only after the downstream splice acceptor, resulting in a shorter 3’UTR region (**Figure 4D, Supplementary Figure 3**). The interaction signal appears to distinguish *MOBP* transcript isoforms with different 3′UTR architecture, rather than reflecting a change in total *MOBP* expression or simple exon inclusion. As 3′UTRs can influence mRNA stability, localisation, translational efficiency and post-transcriptional regulation (35), these findings provide a potential mechanism through which disease-associated genetic variation and local DNA methylation could affect *MOBP* protein output without producing a detectable change in total transcript abundance.

## Discussion

Here, we investigated genetic and epigenetic variation at the *MOBP* locus across multiple neurodegenerative diseases, revealing both shared and disease-specific mechanisms. Analysis of GWAS summary statistics identified significant associations in ALS, PSP, and FTD. Notably, ALS and PSP showed strong overlap in lead risk SNPs and high posterior probabilities of colocalisation, indicating a shared underlying causal variant. Bayesian fine-mapping further supported this convergence, with credible sets in ALS and PSP highlighting a largely overlapping set of variants within the *MOBP* locus. In contrast, FTD demonstrated weak evidence of colocalisation with ALS and PSP, and fine-mapping localised FTD-associated variants to a region upstream of the *MOBP* transcription start site, suggesting a distinct regulatory mechanism. mQTL analyses further highlighted cg15069948 as a key CpG site linking genetic risk to DNA methylation, acting as a significant mQTL for the shared ALS and PSP top risk SNPs. In line with this mQTL effect, in PSP white matter, we also observed lower cg15069948 methylation levels in the risk rs1768208-T. However, we did not observe robust eQTLs or pQTLs linking disease-associated variants to *MOBP* gene expression in available brain datasets. Overall, these findings suggest that *MOBP* eQTL/pQTL variation in oligodendrocytes/brain is unlikely to explain disease risk, and that the underlying genetic mechanisms at this locus are more consistent with mQTL regulation.

There is increasing recognition of the importance of oligodendrocytes and myelin involvement in neurodegeneration (36–41). Many neurodegenerative diseases exhibit well described oligodendrocyte pathology, for example in MSA (42), where they contain characteristic glial cytoplasmic inclusions composed of aggregated α-synuclein. Primary tauopathies such as PSP and corticobasal degeneration (CBD) also exhibit prominent OLG pathology, with tau accumulation forming coiled bodies within these cells (43,44). In ALS, where loss of motor neurons is the major pathological hallmark of disease, there is also growing evidence for oligodendrocyte involvement (45,46). The recurrent identification of myelin-associated genes such as *MOBP* as implicated across multiple neurodegenerative diseases further underscores the importance of oligodendrocyte and myelin biology, highlighting the need to better understand their contribution to disease mechanisms.

Prior epigenetic studies have reported promoter hypermethylation of *MOBP* in MSA, accompanied by reduced gene expression, implicating DNA methylation as a potential regulatory mechanism at this locus (2,17,18). However, we did not observe evidence of genetic association linking *MOBP* to MSA, suggesting that epigenetic dysregulation at this locus in MSA may occur independently of inherited genetic variation. Altered DNA methylation at *MOBP* in MSA provided a strong rationale to investigate whether disease-associated genetic variation at *MOBP* converges on altered DNA methylation across neurodegenerative diseases, and whether such epigenetic changes may reflect shared or disease-specific regulatory pathways. Our findings indicate that while ALS and PSP share a common genetic risk signal at *MOBP* that colocalises with variations at DNA methylation sites, FTD likely involves a distinct genetic and regulatory contribution within the same locus. However, the comparatively smaller FTD and MSA GWAS may have reduced power to detect more modest associations at *MOBP*, and therefore the absence or apparent distinction of genetic signals in these diseases should be interpreted with caution. Together, these results suggest that *MOBP* represents a convergent gene implicated across neurodegenerative diseases, but through different mechanisms, highlighting a nuanced interplay between genetics and epigenetics in disease risk.

The absence of a detectable eQTL at the *MOBP* locus raises the possibility that DNA methylation may influence gene regulation through mechanisms other than steady-state gene expression. DNA methylation within gene bodies and regulatory regions has been implicated in the modulation of transcriptional dynamics, including alternative splicing, transcript processing and exon inclusion or exclusion, rather than overall transcript abundance (47,48). In this context, disease-associated methylation changes at *MOBP* may affect isoform usage, transcript stability, 3′UTR usage or transcriptional elongation in a cell-type-specific manner. Although a limitation of this study is the lack of overlapping PSP/ALS-specific genotype, transcriptomic and DNA methylation data, we observed supportive evidence in an independent dataset that the relationship between cg15069948 methylation and *MOBP* transcript structure differs according to rs1768208 genotype. Specifically, methylation at cg15069948 was associated with usage of an alternative 3’ spice site altering the 3′UTR region in carriers of the PSP/ALS risk allele. These findings suggest that genetic and epigenetic variation at the *MOBP* locus may influence disease risk through effects on transcript processing or post-transcriptional regulation, rather than through detectable changes in total *MOBP* expression. This mechanism may be relevant to our previous observation of altered *MOBP* protein isoform proportions in PSP white matter (2), although further PSP-specific multi-omic data will be required to directly link 3′UTR usage, transcript processing and protein isoform output.

An important question arising from our analyses is why genetic association at the *MOBP* locus appears to cluster across specific neurodegenerative diseases, while being absent or weaker in others. One potential explanation relates to shared underlying pathological processes, particularly the nature of proteinaceous inclusions that define each disease. ALS and PSP show overlapping genetic signals at *MOBP*, and while PSP is classed as a primary tauopathy, ALS, although classically characterised by TDP-43 pathology, has also been associated with altered tau metabolism (49). We also saw a suggestively significant signal at the *MOBP* locus in FTD in a mixed FTD-TDP and FTD-tau cohort (21). Supporting a potential link to tau pathology, the PSP risk allele (T) at rs1768208 has been associated with increased tau thread pathology and phosphorylated tau-positive coiled bodies in oligodendrocytes (15). However, a purely tau associated explanation is unlikely, as we did not see any colocalisation at *MOBP* in AD, despite its classification as a tauopathy. Although *MOBP* has previously been linked to AD, rs1768208 (in high LD with our identified lead SNP at ALS/PSP) was only associated in APOE ε4 positive subjects (13). *APOE*, the strongest genetic risk factor for AD, has recently been implicated in oligodendrocyte dysfunction, with APOE ε4 associated with reduced expression of myelin-related genes and impaired cholesterol handling in oligodendrocytes (50). Together, these observations raise the possibility that genetic variation at *MOBP* may exacerbate oligodendrocyte vulnerability in specific pathological contexts, and that additional factors such as *APOE* genotype may be required for genetic risk at this locus to manifest in AD.

We also considered whether the association was linked to the amount of oligodendrocyte pathology reported in these diseases. As described above, a pathological hallmark of PSP is cytoplasmic inclusions in oligodendrocytes presenting as coiled bodies (13,51) and there is growing evidence as to the involvement of oligodendrocytes in ALS (45,46). In MSA, a disease in which the primary cell type affected by pathology is oligodendrocytes, one would expect that if it was the case that the genetic association was linked to degree of oligodendrocyte pathology, we would see a stronger genetic signal at *MOBP.* Nevertheless, in MSA *MOBP* expression seems to be dysregulated via hypermethylation at the promoter region. Altogether, this suggests that different mechanisms might be affecting the dysregulation of this locus and disease risk differently across diseases, implicating DNA methylation only in MSA and genetic variation influencing DNA methylation in PSP and ALS.

Whilst our analysis into *MOBP* across neurodegenerative diseases has provided important insights, there are some limitations to discuss. As with any analysis utilising publicly available data, we rely upon quality and completeness of datasets. Heterogeneity, co-pathologies and misdiagnosis can confound GWAS analyses, leading to apparent associations that may not be disease-specific while reducing power to detect true genetic effects in specific diseases. We also cannot exclude the possibility of the smaller GWAS being underpowered or brain region specific effects on eQTLs and pQTLs. Additionally, our focus on *MOBP* does not exclude other causal genes at this locus. Recent functional work has implicated the neighbouring gene ribosomal protein SA (*RPSA*) (52), suggesting that associated variants may influence multiple genes within the region. Although regulatory effects on RPSA or other neighbouring genes cannot be completely excluded, and further functional studies would be required to resolve their relative contributions, in our analyses, including an extra 2-Mb region centred on *MOBP* (*MOBP* ±1 Mb), all the prioritised PSP/ALS variants and downstream effects were localised within *MOBP*, further supporting *MOBP* as the primary candidate gene underlying the association at this locus.

## Conclusions

In this study, we integrated genetic, epigenetic and transcriptomic analyses to investigate the role of the *MOBP* locus across neurodegenerative diseases. We identified a shared genetic association signal between PSP and ALS at *MOBP*, which also colocalised strongly with DNA methylation, suggesting that disease-associated variation at this locus may converge on epigenetic regulation. In contrast, the FTD-associated signal appeared genetically distinct, while the previously reported promoter hypermethylation in MSA likely reflects a separate disease-associated mechanism. Together, these findings highlight complex regulatory mechanisms at *MOBP* with both shared and disease-specific contributions to neurodegeneration. More broadly, this work adds to growing evidence that oligodendrocyte biology and myelin-associated pathways are important across multiple neurodegenerative diseases, extending beyond traditional neuron-centred models of pathogenesis. By linking inherited risk at *MOBP* to DNA methylation and altered transcript processing, our findings provide a potential mechanistic bridge between genetic susceptibility and downstream oligodendrocyte dysfunction. Given the dynamic and potentially reversible nature of epigenetic regulation (53), these results also support further investigation of oligodendrocyte-associated epigenetic mechanisms as potential therapeutic targets.

## Declarations

### Ethics approval and consent to participate

This study used secondary analysis of previously published results. All studies from which datasets were utilised had received local ethical approval and informed consent.

### Consent for publication

Not applicable

### Availability of data and materials

#### Code Availability

All code used for analysis is made available in a publicly accessible GitHub repository: https://github.com/CBettencourtLab/MOBPAnalysis.

#### Data Availability

ALS, MSA, AD, CJD and PD GWAS summary statistics were accessed through the GWAS Catalog (accession numbers GCST90027164, GCST90406924, GCST90027158, GCST90001389 and GCST009325, respectively). FTD GWAS summary statistics were accessed from the UCL Research Data Depository (https://doi.org/10.5522/04/25600692.v1.). mQTL, eQTL and pQTL summary statistics are available through the NIAGADS Open Access Data Portal (NG00184). For the matched DNA methylation, genotype and expression/splice junction data, DNA methylation data is available at GSE197305, and genotype data at https://portal.dementiasplatform.uk/.

### Authors’ contributions

KF led all data analysis, prepared the figures, and drafted the manuscript; MM, JH, EP, KL, TR and RdS contributed with datasets and/or interpretation; KWF, JH and CB conceptualised and supervised the study; All authors critically reviewed and approved the manuscript.

## Acknowledgements

We thank Dr Gao Wang and his lab for sharing the methylation QTLs. We would like to thank members of the Raj lab for help in accessing the *BigBrain* data resource. This work was supported in part through the computational resources and staff expertise provided by Scientific Computing at the Icahn School of Medicine at Mount Sinai and supported by the Clinical and Translational Science Awards (CTSA) grant no. UL1TR004419 from the National Center for Advancing Translational Sciences. Research reported in this paper was supported by the Office of Research Infrastructure of the National Institutes of Health under award numbers S10OD026880 and S10OD030463. The content is solely the responsibility of the authors and does not necessarily represent the official views of the National Institutes of Health.

