## Supplementary Fig for "Genetic and epigenetic convergence at *MOBP* links ALS and PSP"

*Corresponding Authors:

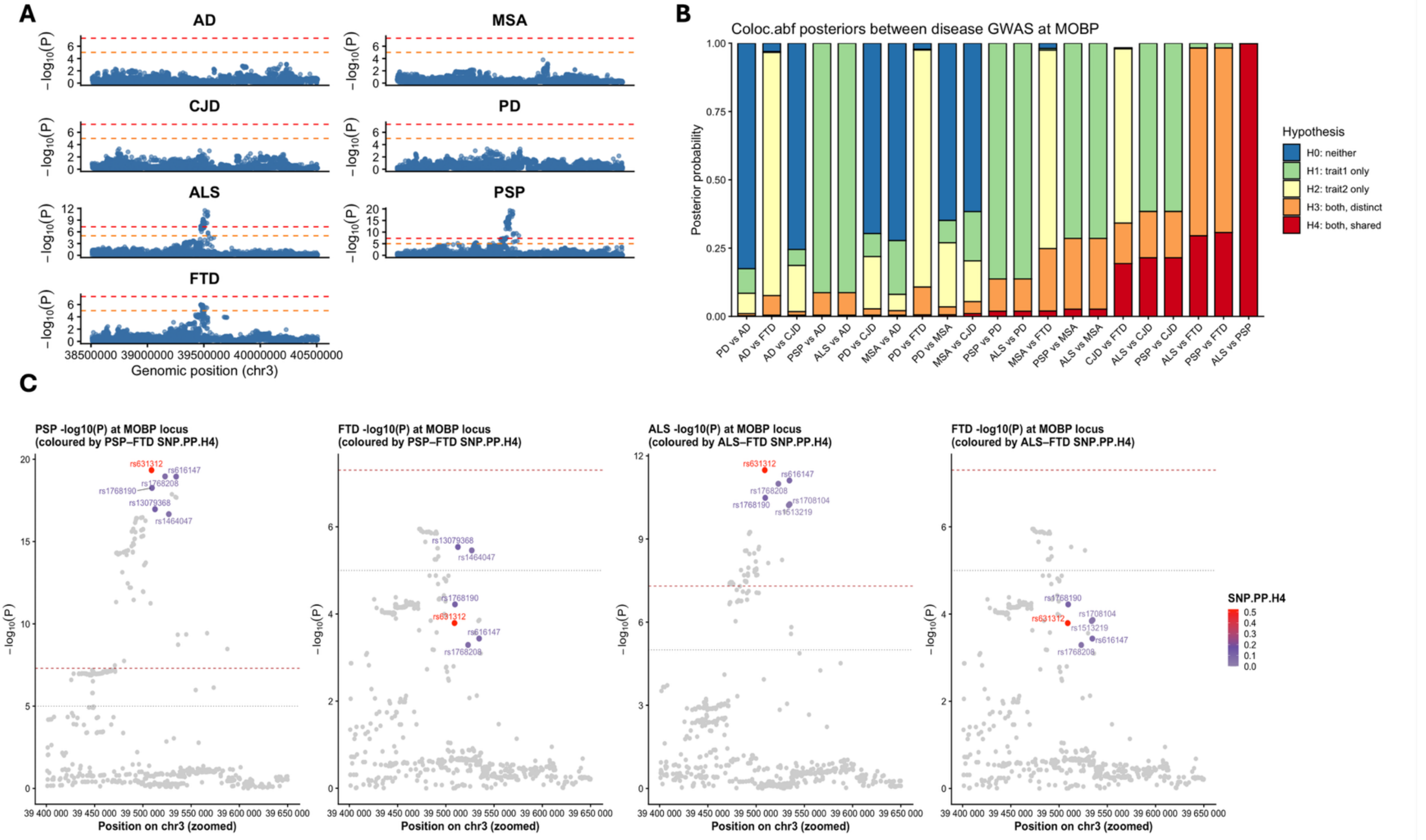


### Supplementary Fig. 1 | Genetic architecture and cross-disease colocalisation of the *MOBP* locus across neurodegenerative diseases.

A) Regional Manhattan plots for AD, MSA, CJD, PD, ALS, PSP and FTD across a ±1 Mb window around the *MOBP* locus (chr3). Points represent single SNP associations, plotted as $-\log_{10}(P)$. Red and orange dashed lines indicate genome-wide significance ($P = 5\times10^{-8}$) and suggestive significance ($P = 1\times10^{-5}$), respectively. B) Pairwise correlation heatmap of GWAS $-\log_{10}(P)$ values across the *MOBP* region. C) COLOC.ABF posterior probability distribution for all pairwise GWAS comparisons at the MOBP locus. Bars show posterior probabilities for hypotheses H0–H4: H0 = neither trait associated; H1 = trait 1 only; H2 = trait 2 only; H3 = both traits associated but with distinct causal variants; H4 = both traits share a causal variant. C)  Zoomed association plots for PSP, ALS and FTD around the *MOBP* peak, with SNPs coloured by their SNP-level posterior probability of belonging to the shared H4 component from COLOC.ABF (SNP.PP.H4). ALS, amyotrophic lateral sclerosis; PSP, progressive supranuclear palsy; FTD, frontotemporal dementia; AD, Alzheimer's disease; MSA, multiple system atrophy; PD, Parkinson's disease; CJD, Creutzfeldt-Jakob disease; SNP, single-nucleotide polymorphism; PP, posterior probability.

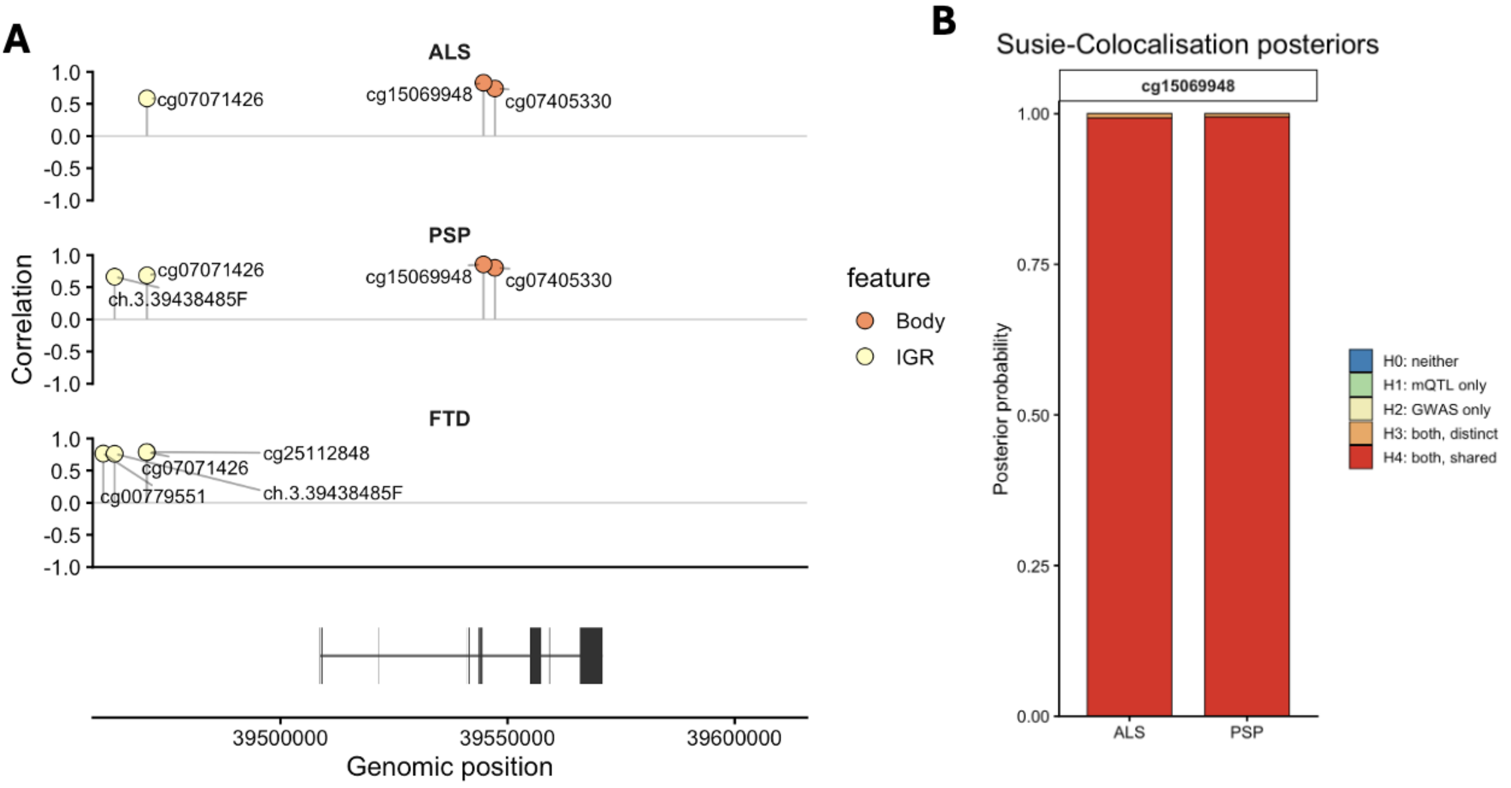


### Supplementary Fig. 2 | mQTLs at *MOBP*

A) Correlation between mQTL and GWAS signals across top CpG sites within the MOBP locus for ALS, PSP, and FTD. Each point represents a CpG site, plotted by genomic position. Height of lollipop point represents Pairwise correlation of GWAS –log10(P) against mQTL -log10(P) values across the MOBP region for ALS, PSP and FTD. CpGs are annotated and coloured by genomic feature. The gene structure for MOBP is shown below.B) Bars show posterior support for each hypothesis: H0 (no association), H1 (disease only), H2 (CpG only), H3 (distinct causal variants), and H4 (shared causal variant). B) Genomic schematic of overlapping CS1 SNPs between ALS and PSP at the *MOBP* locus (blue lollipops) and the cg15069948 CpG site (green lollipop). *MOBP* exons are shown in black, and distance between the lead SNP (rs631312) and the CpG is indicated. ALS, amyotrophic lateral sclerosis; PSP, progressive supranuclear palsy

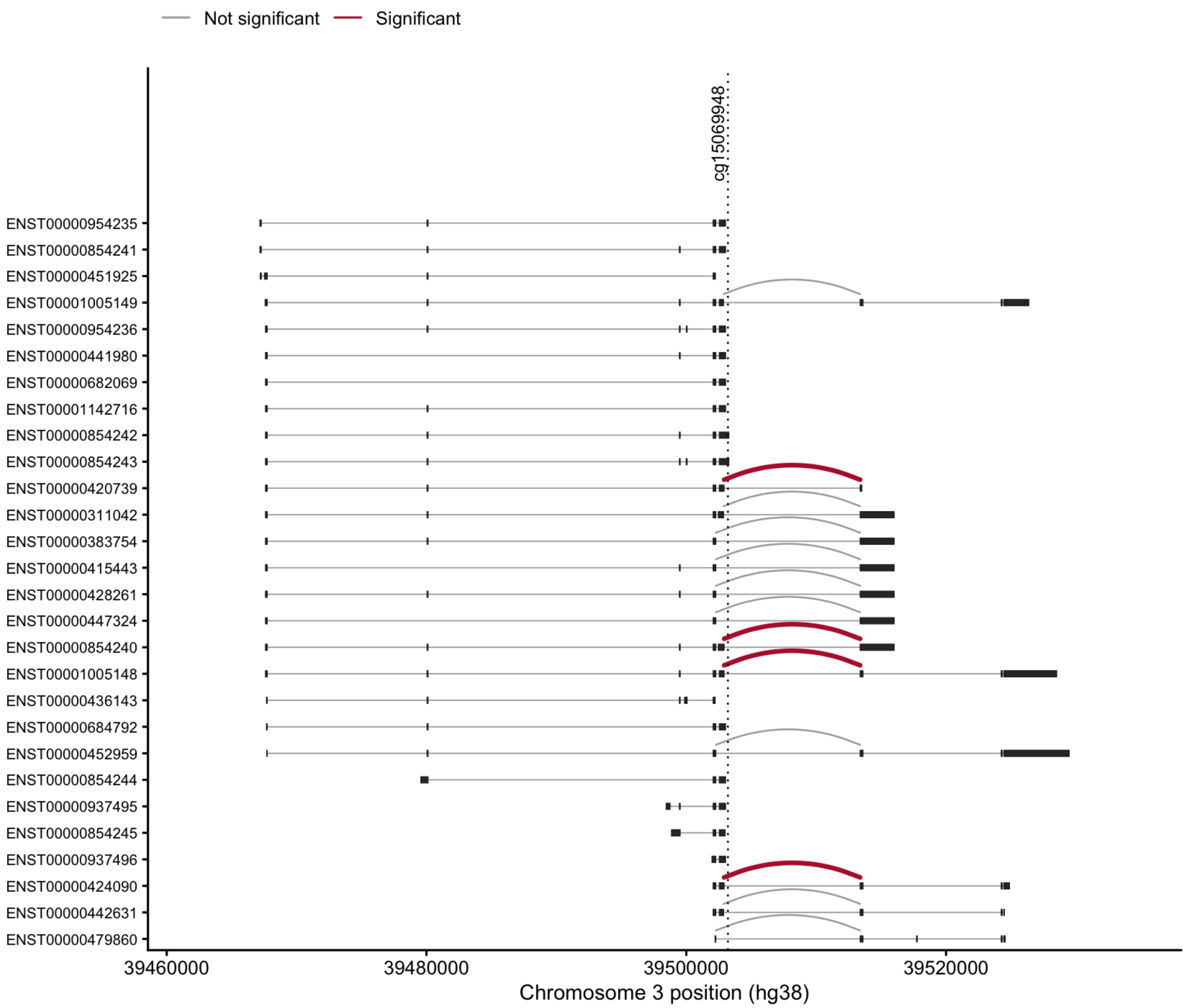


**Supplementary Fig. 3 | Transcript-level context for genotype-by-methylation interaction-associated *MOBP* splice junctions.**

Schematic representation of annotated *MOBP* transcript structures across the cluster 890 region, highlighting splice junctions tested for interaction between rs1768208 genotype and cg15069948 DNA methylation. Each horizontal line represents an annotated *MOBP* transcript, with exon boundaries indicated by black blocks/vertical marks. Arcs represent splice junctions leaving exon 4 and are coloured according to interaction significance, with red arcs indicating junctions showing nominal evidence of genotype-by-methylation interaction and grey arcs indicating non-significant junctions. The position of cg15069948 is shown by the vertical dotted line.
